# Temporal Dynamics of Kidney Mitochondrial Dysfunction in Type 2 Diabetes: Analysis of the Goto-Kakizaki Model

**DOI:** 10.64898/2025.12.27.696702

**Authors:** Bianca Castino, Luciana Jorge, Alef A. C. Santos, André L.L. Bachi, Fernanda T. Borges

## Abstract

Type 2 diabetes mellitus (T2DM) is a major cause of diabetic kidney disease (DKD), a complication driven by chronic hyperglycemia, oxidative stress, and inflammation. Mitochondrial dysfunction has been recognized as a central feature of DKD, but the temporal relationship between mitochondrial alterations, mitophagy, and inflammatory responses remains unclear. Using the non-obese Goto-Kakizaki (GK) rat, a spontaneous model of T2DM, we examined renal mitochondrial changes at 21, 60, and 120 days. Histology revealed a trend toward increased glycation product deposition, more evident at 60 days. Cytochrome C and OPA-1 expression indicated early mitochondrial adaptation through enhanced respiratory activity and hyperfusion, while reduced PGC1-α and PINK suggested impaired biogenesis and mitophagy. By 120 days, PGC1-α and PINK increased, likely reflecting delayed, insufficient compensation, coinciding with mitochondrial depolarization. Cytokine analysis revealed a biphasic inflammatory profile: IL-18 and CXCL1 were sharply elevated at 21 days, IL-1β remained stable, and IL-10 was elevated at 21 and 60 days, indicating early pro-inflammatory activation followed by anti-inflammatory modulation. These findings suggest that renal mitochondrial dysfunction in GK rats evolves from an adaptive state to late-stage failure, with inflammation occurring as transient peaks rather than a persistent process. The period between 21 and 60 days emerges as a critical therapeutic window in which strategies to preserve mitochondrial quality control and modulate inflammasome-related responses may be most effective.

## Introduction

Type 2 diabetes mellitus (T2DM) is a multifactorial metabolic disease characterized by insulin resistance and chronic hyperglycemia, affecting more than 500 million people worldwide and closely associated with complications such as diabetic kidney disease (DKD) (1). Mitochondrial dysfunction has emerged as a central axis in the pathogenesis of T2DM, as impaired mitochondria fail to meet the energy demands of metabolically active tissues such as the kidney, leading to the accumulation of reactive oxygen species (ROS) and activation of inflammatory pathways (2). In the renal proximal tubule, where 90% of the filtered glucose is reabsorbed, hyperactivity of SGLT2 transporters under hyperglycemic conditions exacerbates ATP demand, overloading oxidative phosphorylation and triggering adaptive responses that may progress to irreversible cellular damage (3).

The Goto-Kakizaki (GK) rat, developed in the 1970s through selective breeding of Wistar rats with glucose intolerance, is a unique non-obese T2DM model that spontaneously develops insulin resistance and progressive hyperglycemia without pharmacological or dietary interventions (4). The absence of confounding variables such as obesity and lipotoxicity allows the isolation of chronic hyperglycemia’s effects on renal pathophysiology, making the GK model an ideal tool for studying early stages of diabetic kidney disease (5). However, the timeline of mitochondrial alterations in the kidneys of this model remains poorly explored, particularly regarding the interplay between biogenesis, mitophagy, and inflammation.

Mitochondria are dynamic organelles essential for cellular homeostasis, functioning not only as energy centers via oxidative phosphorylation but also regulating processes such as apoptosis, redox balance, and calcium signaling (6). Under physiological conditions, mitochondrial dynamics—fusion (organelle integration) and fission (fragmentation)—maintain a functional network, allowing for damage repair and the balanced distribution of essential components (7). Fusion, mediated by proteins such as OPA-1, facilitates the complementation of mitochondrial DNA (mtDNA), while fission, regulated by DRP1, isolates damaged mitochondria for degradation via mitophagy (a process dependent on PINK1/PARKIN) (8). In disease states such as T2DM, chronic hyperglycemia and oxidative stress disrupt mitochondrial balance, impairing not only ATP production but also the ability to adapt to metabolic fluctuations (9). The dysregulation of mitochondrial dynamics, crucial for maintaining a functional network, creates a favorable environment for the accumulation of macromolecular damage, which, in highly energy-dependent tissues like the kidney, accelerates the loss of cellular function (10). This dysfunction, combined with the inability to remove damaged components, establishes a vicious cycle of redox stress and inflammatory signaling—key factors in the pathogenesis of complications such as diabetic kidney disease (11).

Diabetic kidney disease (DKD), present in approximately 40% of patients with T2DM, is driven by complex mechanisms including glomerular hyperfiltration, oxidative stress, and the accumulation of advanced glycation end products (AGEs) (12). Recent studies highlight that mitochondrial dysfunction precedes structural alterations such as basement membrane thickening, suggesting that early interventions targeting energy homeostasis could slow disease progression (13). However, most research focuses on advanced stages of DKD, neglecting the critical transition between compensatory adaptation and metabolic failure.

This study aimed to investigate the temporal progression of renal mitochondrial dysfunction in the Goto-Kakizaki model of type 2 diabetes, identifying critical transition points between metabolic adaptation and cellular failure, with an emphasis on the relationship between mitochondrial dynamics, mitophagy, and inflammation.

## Materials and Methods

### Ethics Committee

This study was conducted in accordance with the guidelines established by the Ethics Committee on Animal Use (CEUA) of the Federal University of São Paulo (UNIFESP), under approval number 3152021019. All experiments complied with current regulations to ensure animal welfare and adherence to ethical standards.

### Experimental Groups

Renal tissues used in this study were provided by the Institute of Physical Activity and Sports Sciences (ICAFE), Universidade Cruzeiro do Sul. Samples were obtained from Goto-Kakizaki and Wistar rats at three ages: 21, 60, and 120 days, forming the GK21D, GK60D, and GK120D groups, with their respective controls W21D, W60D, and W120D. Each group consisted of at least four animals and each protocol was repeated twice.

### Model Characterization

The diabetic status of the animals was previously characterized by our research group and described in earlier studies (14–18). In Gimenes et al., 2024, fasting blood glucose was assessed five days before euthanasia, following a 12-hour fasting period, using blood collected from the tail and measured with a glucose monitor (AccuCheck, Corydon, IN, USA). Results showed that 120-day-old GK rats had significantly higher fasting glucose levels compared to Wistar controls (159.3 ± 5 vs. 78.2 ± 5.5), confirming the diabetic condition of the colony. Additionally, body weight analysis revealed that diabetic rats exhibited lower body mass (347.1 ± 3.7 vs. 508.4 ± 11) compared to the control group, consistent with the Goto-Kakizaki model’s pathophysiology.

### Histology

Kidneys were aseptically removed, decapsulated, longitudinally sectioned, washed with saline, and fixed in 10% buffered formalin (pH 7.4). After fixation, tissues were processed in the Pathology Laboratory, embedded in paraffin, and sectioned using a microtome for histological slide preparation.

### Periodic Acid-Schiff (P.A.S) Staining

Slides were deparaffinized in xylene, rehydrated in alcohol gradients, and washed rinsed in distilled water. They were then incubated with periodic acid (10 min), washed in distilled water (3 min), and treated with Schiff reagent (15 min, dark chamber), followed by another wash (10 min). Counterstaining was performed with Carazzi’s hematoxylin (4 min), and slides were dehydrated, mounted, and analyzed under a light microscope (LEICA), with quantification expressed as %/area.

### Immunohistochemistry

Slides were deparaffinized, rehydrated, and subjected to antigen retrieval (95°C, Tris/EDTA 1 mmol/L, pH 9.0, 30 min). Endogenous peroxidase activity was blocked with 5% H2O2 (10 min, room temperature). Primary antibody incubation followed (18 h, 4°C), with subsequent washing in TBS and incubation with HRP-conjugated secondary antibody (1 h, LSAB Kit, DAKO). Slides were counterstained with hematoxylin, dehydrated, and analyzed under a light microscope (LEICA), with quantification expressed as %/area. Negative controls omitted the primary antibody.

Primary antibodies used: PGC1-alpha (1:500, Cell Signaling #6946); Acetyl-CoA (1:500, Cell Signaling #3676); GLS-2 (1:500, Abcam #113509); Cytochrome C oxidase (1:500, Abcam #90529).

### Western Blotting

Kidneys were homogenized in Radio-Immunoprecipitation Assay buffer (RIPA) with protease inhibitors for protein extraction. Protein concentration was determined using the Lowry method with BSA standards and spectrophotometric reading (595 nm). Protein separation was performed in an acrylamide gel (30% acrylamide, 1M tris-HCl, 10% SDS) with stacking and resolving gels. Electrophoresis was conducted in a vertical chamber (Mini-Protean II, Bio-Rad, USA) at an initial voltage of 60V, followed by 120V (90 min). Protein transfer was carried out on a nitrocellulose membrane (Trans-Blot® Turbo™ RTA, Bio-Rad, USA) activated in cold buffer (5× Trans-Blot® Turbo™ Buffer, 100% ethanol). After assembling the transfer “sandwich,” the process was performed using the Trans-Blot® Turbo™ Transfer System (Bio-Rad, USA) for 20 min (25V-1.0A-30M). The membrane was kept in TBS-T before immunodetection. For detection, the membrane was blocked with 5% BSA in TBS-T (1 h), washed, and incubated with the primary antibody overnight. After further washes, incubation with the secondary antibody followed (1 h), followed by revelation with ECL Plus (GE Healthcare, Russia) and capture using the Amersham™ Imager 600 (Amersham Biosciences, Germany). Band intensity was automatically quantified using ImageJ software and expressed as the ratio of each protein normalized to β-actin or GAPDH.

### Determination of cytokine levels in renal tissue by ELISA

Relative levels of CXCL1, IL-10, IL-1β, and IL-18 in renal tissue were assessed by sandwich ELISA using the DuoSet® ELISA Development System (R&D Systems, Minneapolis, MN, USA; catalog numbers DY275, DY217B, DY201, and DY318). Kidney fragments were homogenized in PBS (pH 7.4) at 1:10 (w/v), centrifuged at 12,000 × g for 15 min at 4 °C, and the supernatants used immediately for ELISA and total protein quantification by Bradford assay. High-binding plates (Nunc MaxiSorp™, Thermo Fisher Scientific) were coated overnight at 4°C with capture antibodies diluted in carbonate-bicarbonate buffer (0.05 M, pH 9.6). After blocking with 1% BSA for 1 h at room temperature, samples were added in duplicate, followed by biotinylated detection antibodies and streptavidin-HRP. The reaction was developed with TMB, stopped with 2 N H2SO_4_, and absorbance read at 450 nm with 570 nm correction (BioTek®).

Data were normalized and expressed as a percentage relative to the control group mean (100%).

### Isolation and Culture of Proximal Tubule Cells

Kidneys from 120-day-old GK and Wistar rats were extracted and immersed in PBS 1X + 1% P/S for decapsulation. Tissues were minced (~1 mm^3^) and incubated (37°C, 30 min, 300 rpm) in a digestion solution (PBS 1X + 0.3 mL of type I collagenase at 25 mg/mL). After digestion, cells were homogenized using a 3-mL syringe with a 20G needle until resistance was absent, filtered (70 µm sieve), and centrifuged (300 × g, 5 min, 4°C). The pellet was resuspended in 2 mL of PBS 1X and subjected to Percoll 28% separation (2 mL of cell suspension in 10 mL of Percoll 28%), followed by centrifugation (2200 rpm, 10 min, 4°C). The pellet containing proximal tubule cells was washed twice with PBS 1X and centrifuged (300 × g, 5 min). Cells were then resuspended in 4 mL of Renal Epithelial Cell Medium-2 supplemented with SupplementMix and 1% P/S, plated in 10 cm culture dishes (2 mL/dish), and incubated (37°C, 5% CO_2_). The medium was changed the next day. Once 80% confluence was reached, cells were trypsinized (0.25% trypsin-EDTA), counted, and seeded in black 96-well plates (10^5^ cells/well), incubated (37°C, 5% CO_2_) for analysis in the Incell Analyzer 2000.

### Mitotracker and MitoSOX™ Red

Mitotracker® Red CMXRos (Thermo Fisher, M7512) was diluted in DMSO for a 1 mM stock solution and further diluted in serum-free medium to 200 nM. After medium removal, the solution was added to the cells and incubated for 30 min (37°C, protected from light). Cells were then washed twice with PBS (pH 7.4) and incubated with DAPI (1 µg/mL in PBS, 10 min, room temperature, protected from light). After another PBS wash, fluorescence was captured using the Incell Analyzer 2000 (excitation/emission: Mitotracker 579/599 nm; DAPI 358/461 nm). For MitoSOX™ Red (Thermo Fisher, M36008), the stock solution (5 mM in DMSO) was diluted in PBS to 5 µM. After medium removal, the solution (100 µL/well) was added to the cells and incubated for 10 min (37°C, protected from light). After two PBS washes, cells were incubated with DAPI (1 µg/mL, 10 min, room temperature, protected from light), washed, and analyzed using the Incell

Analyzer 2000 (excitation/emission: MitoSOX™ Red 510/580 nm; DAPI 358/461 nm). Data were processed using the associated software for fluorescence intensity quantification and nuclear staining visualization.

### Statistical Analysis

Statistical analysis was performed using GraphPad Prism 7.0.

## Results

PAS staining revealed a tendency for the accumulation of glycation end-products in the renal tissues of GK animals (Figure 1, A-B), indicative of AGEs. At 21 days, the GK group exhibited 22 ± 1.7% of stained area, compared to 20.4 ± 1.5% in the Wistar group; at 60 days, the values were 24.7 ± 1.9% (GK) versus 19.04 ± 1.2% (Wistar); and at 120 days, 21.6 ± 1.8% in GK compared to 20.5 ± 1.4% in Wistar. Although not statistically significant, this trend suggests a possible progressive accumulation, particularly at 60 days.

**Figure 1.**
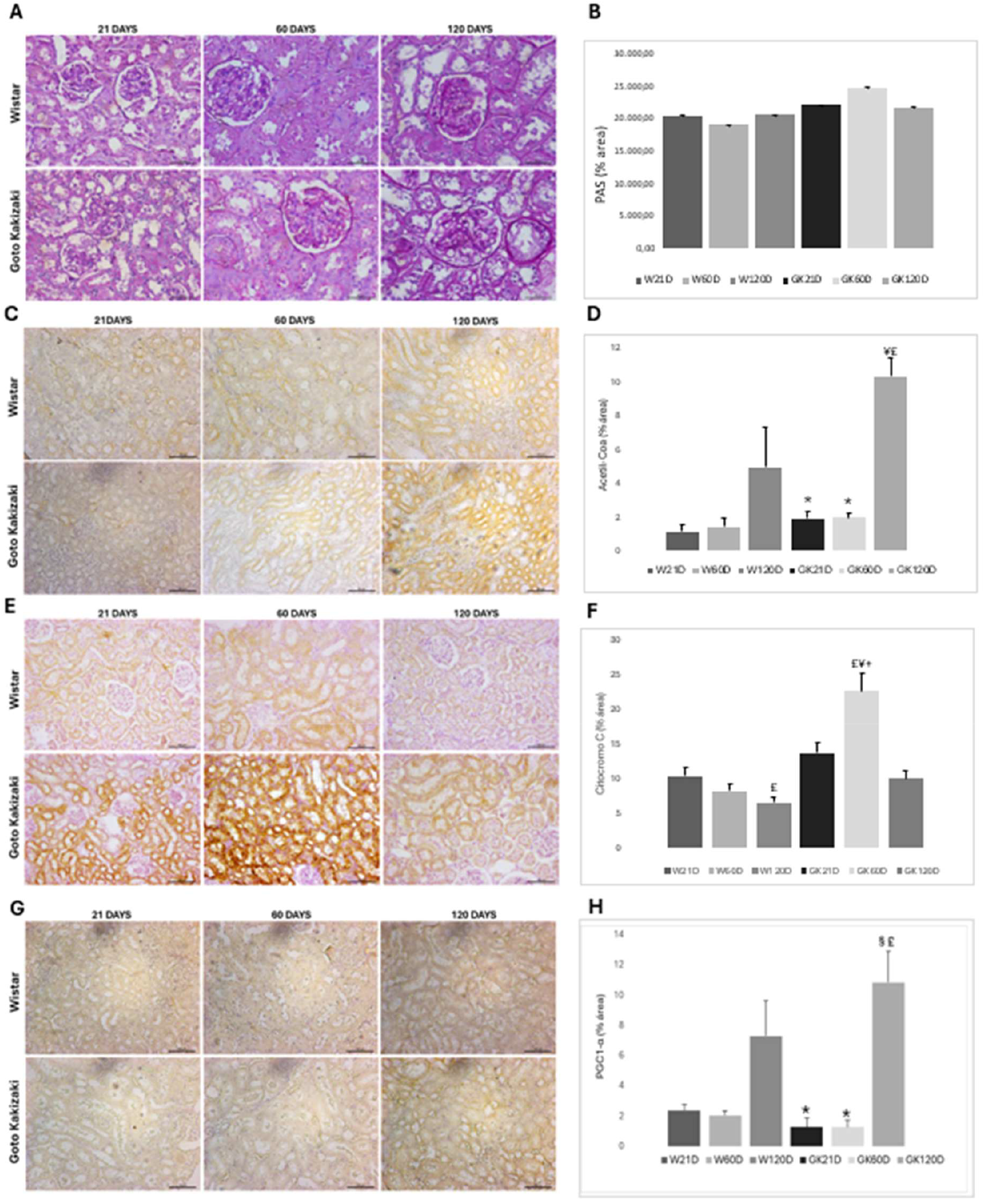
Histological and bioenergetic alterations in the kidneys of GK and Wistar rats over time. (A-B) Periodic acid-Schiff (PAS) staining in renal sections, highlighting progressive structural changes in the GK group and quantification of PAS-stained area (% of total area). (C-D) Immunohistochemistry for acetyl-CoA and its quantification. (E-F) Immunohistochemistry for cytochrome C oxidase and its quantification. (G-H) Immunohistochemistry for PGC-1α and its quantification, indicating variations in mitochondrial biogenesis associated with the progression of the diabetic phenotype. Data are expressed as mean ± standard deviation. *p < 0.05 vs. Wistar group at the same experimental time; § p < 0.05 vs. GK 21D; £ p < 0.05 vs. GK 60D; £¥+ indicates statistically significant differences between groups and time points analyzed.

In the same context, the expression of Acetyl-CoA (Figure 1, C-D) in renal tissue reinforces the presence of metabolic alterations. From the earliest time points, the GK group exhibited a greater stained area (1.93 ± 0.4% at 21 days) compared to Data were expressed as Mean ± Standard Error of the Mean (SEM), with normality assessed using the Shapiro-Wilk test. Group comparisons were conducted using one-way ANOVA followed by Tukey’s test. Comparisons between two groups were performed using Student’s t-test. The significance level was set at p < 0.05.

Wistar (1.15 ± 0.4%), maintaining this difference at 60 days (1.94 ± 0.4% vs. 1.44 ± 0.3%). At 120 days, the increase was even more pronounced in GK (10.28 ± 2.3%) compared to Wistar (4.96 ± 1.1%).

Mitochondrial function assessment also revealed important differences. Immunohistochemistry for Cytochrome C (Figure 1, E-F) showed that at 21 days, the GK group had 13.58 ± 1.53% of stained area, higher than the 10.21 ± 1.19% observed in Wistar. This value increased at 60 days, reaching 22.56 ± 2.55% in GK, versus 8.10 ± 1.06% in Wistar. At 120 days, there was a reduction, with 9.92 ± 1.08% in GK and 6.44 ± 0.74% in Wistar.

Similarly, the expression of PGC1-α (Figure 1, G-H), a key protein in mitochondrial biogenesis regulation, was lower in GK at early time points (1.27 ± 0.40% vs. 2.39 ± 0.62% at 21 days and 1.26 ± 0.29% vs. 2.04 ± 0.48% at 60 days), increasing significantly at 120 days (10.86 ± 2.31% in GK versus 7.31 ± 2.03% in Wistar). These data suggest an initial impairment in mitochondrial adaptation capacity, followed by a late compensatory response.

Regarding mitochondrial dynamics in kidney cortex, OPA-1 expression, a regulator of mitochondrial fusion, (Figure 2, A-B) was higher in the GK group from day 21 (0.69 ± 0.01 in GK vs. 0.35 ± 0.01 in Wistar), remained similar at day 60 (0.68 ± 0.01 vs. 0.37 ± 0.01), and converged at day 120 (1.07 ± 0.01 in GK and 1.08 ± 0.01 in Wistar). This finding suggests an initial compensatory response hyperglycemic stress.

**Figure 2.**
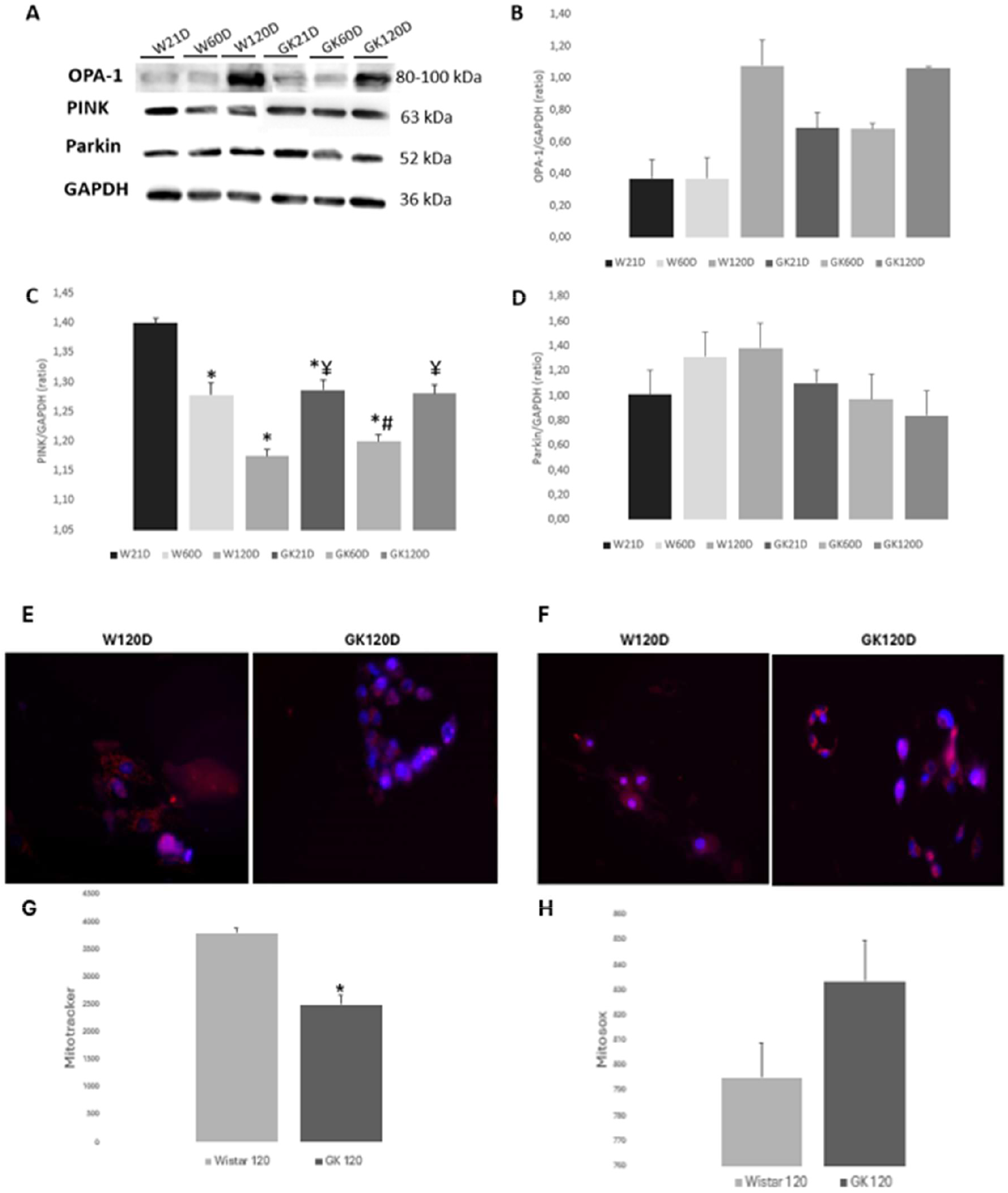
Evaluation of mitochondrial dynamics and oxidative stress. (A-B) Protein expression analysis through Western Blotting and its respective graphical representation in kidney cortex; (A-C) PINK; (A-D) Parkin. (E-G) Mitotracker in isolated proximal tubule cells and its graphical representation. (F-H) Mitosox in proximal tubule cells and its graphical representation. Data are presented as mean ± standard error of the mean (SEM). P<0.05, with (*) 05 vs. Wistar 21D; (#) vs. Wistar 60D; (¥) p < 0.05 vs. Wistar 120D.

Mitophagy assessment revealed divergent patterns between groups. PINK expression (Figure 2, A-C) was lower in GK at the early time points (1.30 ± 0.02 at day 21 vs. 1.40 ± 0.02 in Wistar; 1.18 ± 0.02 at day 60), whereas at day 120, GK rats exhibited higher expression (1.28 ± 0.02 vs. 1.17 ± 0.02). Conversely, PARKIN protein (Figure 2, A-D) was elevated in GK at day 21 (1.10 ± 0.03) but decreased at days 60 (0.77 ± 0.02) and 120 (0.84 ± 0.02), while Wistar animals showed a progressive increase (1.01 ± 0.02 at day 21; 1.32 ± 0.02 at day 60; 1.39 ± 0.02 at day 120).

Furthermore, mitochondrial functionality in proximal tubular cells was assessed through Mitotracker uptake (Figure 2, E-G) and reactive oxygen species (ROS) production via MitoSOX (Figure 2, F-H). At day 120, to the GK group exhibited lower Mitotracker uptake (2494 ± 162 fluorescence units) compared to Wistar (3791 ± 95 fluorescence units), suggesting impaired mitochondrial membrane potential. MitoSOX measurement revealed slightly higher ROS production in GK (836 ± 16 fluorescence units) than in Wistar (795 ± 14 fluorescence units), although this difference was not statistically significant.

ELISA analysis revealed distinct cytokine profiles in the kidney cortex of GK and Wistar rats across different time points (Figure 3, A-D). The analysis of IL1-β (3-A) in GK rats revealed varying levels at different time points. The GK 21 group showed a tendency towards slightly lower cytokine levels, at 95% of the control (SE = 9.3). In contrast, the GK 60 group demonstrated a small increase, reaching 117.7% of the control (SE = 7.6). The GK 120 group, however, exhibited a more notable reduction, with levels dropping to 71.1% of the control (SE = 5.1). For IL-18 (Figure 3-B), a more drastic change was observed. The GK 21 group presented a significant increase, with cytokine levels reaching 284% of the control (SE = 18). On the other hand, the GK 60 and GK 120 groups showed substantial reductions, reaching 57.7% (SE = 10.4) and 66.6% (SE = 8.5) of the control, respectively. In the analysis of CXCL-1 (Figure 3-C), the GK 21 group exhibited a 57.9% increase significant compared to the control, with cytokine levels corresponding to 157.9% of the control (SE = 16). The GK 60 group showed levels close to the control, at 92.2% (SE = 15.5), while the GK 120 group demonstrated a considerable reduction, with only 63.7% of the control (SE = 3.7). The results of IL-10 (Figure 3-D) indicated a persistent increase in the earlier groups. The GK 21 group had an 84.2% increase relative to the control, reaching 184.2% (SE = 15.7). The GK 60 group maintained this upward trend, with levels at 175.6% of the control (SE = 16.2). The GK 120 group, however, showed a normalization of cytokine levels, reaching 87.4% of the control (SE = 7.1).

**Figure 3.**
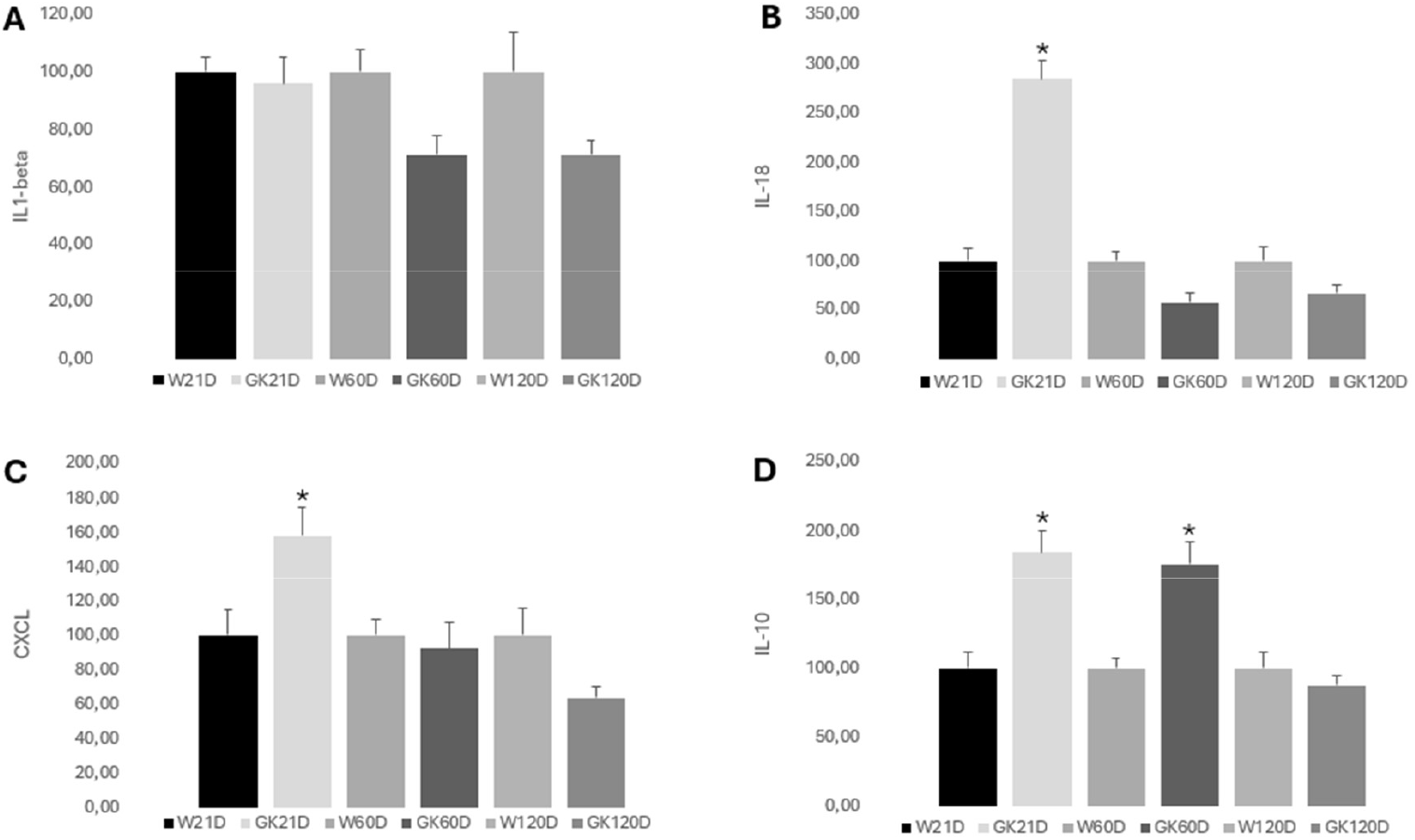
Cytokine levels in the kidneys of GK and Wistar rats over time assessed by ELISA. **(A)** IL-1β levels showing no significant differences across the groups. **(B)** IL-18 levels with a significant increase in the GK21D group compared to all other groups. **(C)** CXCL1 levels with a significant increase in the GK21D group compared to all other groups. **(D)** IL-10 levels showing significantly elevated concentrations in the GK21D and GK60D groups compared to the GK120D group. Data are expressed as percentage relative to the control group and presented as mean ± standard error of the mean (SEM).

## Discussion

Diabetic kidney disease (DKD) is a severe complication of type 2 diabetes mellitus (T2DM), affecting approximately 40% of patients and being one of the leading causes of end-stage renal disease globally (19). Characterized by glomerular hyperfiltration, thickening of the basement membrane, and progressive loss of renal function, its pathogenesis is complex, involving oxidative stress, chronic inflammation, and the accumulation of advanced glycation end products (AGEs) (20). In recent years, mitochondrial dysfunction has been recognized as a central factor in this process, as compromised mitochondria fail to meet the energy demands of metabolically active tissues, such as the kidney, promoting the accumulation of reactive oxygen species (ROS) and the activation of inflammatory pathways (21). In this study, we used the Goto-Kakizaki (GK) model of T2DM, a non-obese model that develops spontaneous hyperglycemia, to investigate the temporal progression of renal mitochondrial dysfunction and its interrelationship with mitochondrial dynamics, mitophagy, and inflammation. Our findings reveal a sequence of metabolic, mitochondrial, and inflammatory changes that evolve over time, highlighting a critical transition between compensatory adaptation and cellular failure.

The analysis of the metabolic profile revealed a tendency for the accumulation of glycation products in the kidney tissues of GK animals, as evidenced by PAS staining. At 60 days, although the differences did not reach statistical significance, this pattern suggests that even in the early stages, chronic hyperglycemia favors the formation of AGEs, known to induce oxidative stress and inflammation through the activation of the RAGE receptor (22). The lack of significance may be related to interindividual variability or the early stage of the disease during the analyzed periods. In parallel, the expression of Acetyl-CoA showed a significant increase in the GK, particularly at 120 days, indicating greater metabolic activity. This increase may represent a compensatory response to the high energy demand imposed by SGLT2-mediated tubular glucose reabsorption, a process that overloads oxidative phosphorylation in the proximal tubule (23). The accumulation of Acetyl-CoA, a central metabolite in the Krebs cycle, may also indicate a change in subsequent metabolic utilization, consistent with cellular stress observed in hyperglycemic conditions (24).

The evaluation of mitochondrial function revealed variations over time. Immunohistochemistry for Cytochrome C indicated an initial increase in the GK at 21 days compared to Wistar, followed by a reduction at 120 days. This pattern suggests an initial adaptive response, with higher activity of the respiratory chain to meet the elevated energy demand, followed by mitochondrial failure in later stages. This transition is consistent with studies linking chronic oxidative stress to the progressive loss of respiratory chain components in T2DM models (25). The expression of PGC1-α, an essential regulator of mitochondrial biogenesis, showed a reduction in the GK at 21 and 60 days, suggesting early impairment in mitochondrial renewal. Notably, at 120 days, there was a marked increase in this marker, possibly reflecting a late compensatory attempt. However, given the already established dysfunction, this response may be insufficient to reverse the accumulated damage (26).

Mitochondrial dynamics also showed significant modifications. The expression of OPA-1, a regulator of mitochondrial fusion, was elevated in the GK animals from 21 days, remaining high until 60 days, before normalizing at 120 days. This pattern suggests an initial phase of hyperfusion as a compensatory mechanism, facilitating the dilution of damaged components and the preservation of mtDNA (27). However, the late normalization may indicate that this adaptation is temporary and becomes insufficient in the face of persistent hyperglycemic stress. Regarding mitophagy, the expression of PINK was reduced in the GK at early stages, suggesting a compromised ability to eliminate dysfunctional mitochondria, which may favor the accumulation of damaged organelles (28).

The inflammatory profile in the kidney of GK model revealed an early, transient activation pattern, rather than sustained high-grade inflammation. The marked increase in IL-18 at 21 days, together with the context of mitochondrial dysfunction and oxidative stress, is compatible with acute activation of the NLRP3 inflammasome. In contrast, IL-1β levels remained relatively stable over time, suggesting that this cytokine may not have been persistently processed or released, possibly due to early activation control mechanisms. The significant elevation of IL-10 at 21-and 60-day points to the engagement of anti-inflammatory pathways capable of modulating the initial pro-inflammatory burst. This pattern supports a biphasic inflammatory response: an acute peak, likely triggered by metabolic stress and mitochondrial damage, followed by a regulatory phase that limits sustained inflammasome activity. While this modulation may protect against prolonged inflammation, it occurs alongside ongoing mitochondrial dysfunction, which may predispose the kidney to recurrent low-grade inflammatory episodes in later stages of disease progression. This interaction underscores the close relationship between mitochondrial homeostasis and the inflammatory response in the progression of renal impairment.

Our results indicate that renal mitochondrial dysfunction in the GK model of T2DM follows a dynamic pattern, with an initial adaptive response—evidenced by increased mitochondrial fusion, respiratory chain activity, and attempts at mitophagy— that evolves into progressive failure in later stages. Mitophagy dysregulation and the interplay between acute inflammasome activation and anti-inflammatory modulation emerge as central factors in the intensification of renal damage, highlighting the need for early interventions to restore mitochondrial homeostasis and balance inflammatory responses. These findings emphasize a critical time window in this model, such as the initial adaptation period at 60 days, during which therapeutic strategies could be more effective. Future studies should investigate approaches such as the activation of PGC1-α to stimulate mitochondrial biogenesis, the use of compounds that promote selective mitophagy, and the modulation of specific inflammatory pathways, including targeted inhibition of NLRP3 (35,36).

## Conclusion

In the Goto-Kakizaki model, insulin resistance and hyperglycemia promote early renal mitochondrial adaptations in response to the excessive energy demand from tubular glucose reabsorption. However, by 60 days, mitochondrial dysfunction reaches a critical threshold: the inefficiency of the mitophagy pathway leads to the accumulation of damaged organelles, oxidative stress, and inflammation, even in the absence of histological alterations. This period emerges as a therapeutic window, where interventions targeting mitochondrial autophagy and redox balance could interrupt the cascade of metabolic damage, offering a promising strategy to halt the development of diabetic kidney disease before irreversible complications are established.

## Author Contributions

Writing - Original Draft Investigation, BC, LJ; Validation, LJ, FTB; Methodology, BC, LJ, AACS and ALLB; Writing-Reviewing and Editing, BC, LJ and FTB; Formal analysis; Visualization, LJ and FTB; Resources, ALLB and FTB; Funding acquisition, FTB; Project administration, FTB; Conceptualization, FTB; Supervision, FTB. All authors have read and agreed to the published version of the manuscript.

## Acknowledgments

Conselho Nacional Científico de Desenvolvimento Tecnológico (CNPq), Universidade Estudos e Projetos (FINEP), Fundação Oswaldo Ramos (FOR), Fundação de Amparo à Pesquisa do Estado de São Paulo (FAPESP), and Coordenação de Aperfeiçoamento de Pessoal de Nível Superior (CAPES).

